# Evolution of virus and virophage facilitates persistence in a tripartite microbial system

**DOI:** 10.1101/2023.01.31.526414

**Authors:** Ana del Arco, Matthias G. Fischer, Lutz Becks

## Abstract

Tripartite biotic interactions are inherently complex, and the strong interdependence of species and high levels of exploitation can make these systems short-lived and vulnerable to extinction. The persistence of species depends then on the balance between exploitation and avoidance of exploitation of the resource beyond the point where sustainable exploitation is no longer possible. We used this general prediction to test the potential for long-term persistence in a recently discovered tripartite microbial system in which a eukaryotic host is preyed upon by a giant virus that is in turn parasitized by a virophage. Host and virophage may benefit from this interaction because the virophage reduces the harmful effects of the giant virus on the host population over time and the virophage can survive integrated into the host genome when giant viruses are scarce. Here, we grew hosts in the presence and absence of the giant virus and virophage over ~280 host generations. We found that the three players persisted, but that the beneficial effect of the virophage for the host population diminished over time. We further tested whether the level of exploitation and replication evolved in the giant virus and/or virophage population over the course of the experiment and whether the changes were such that they avoid overexploitation. We found that the giant virus evolved towards lower replication levels and the virophage towards increased replication but decreased giant virus exploitation. These changes are predicted to facilitate persistence by lowering giant virus and host exploitation and consequently reducing the protective effect of the virophage.

## Introduction

Virophages are a distinct class (*Maveriviricetes*) of viral agents that parasitize the cytoplasmic factories of giant viruses in the order *Imitervirales* for their replication (1). Metagenome studies found that virophages can be highly abundant and diverse, which suggests that they can play an important role for the dynamics of microbial communities through their effects on host-virus dynamics (2). The interaction between hosts and virophages has been described as being mutualistic because virophages facilitate host persistence in the presence of high giant virus densities, and because host cells provide an opportunity for the virophage to persist integrated into the genome of the host when giant viruses are scarce (3–6). The benefit for the host arises because virophages reduce the lytic impact of the giant virus on the host population over time by inhibiting giant virus replication. However, there are few data demonstrating long-term two-way interactions between host and virophage at the population level. Like other tripartite systems with strong interdependence of interacting species and high levels of exploitation (e.g., hyperparasitism, satellite viruses and helper viruses (48, 49)), host-virus-virophage systems may be short-lived and prone to extinction (e.g. 5, 6).

Long-term persistence of host, giant virus and virophage depends largely on the severity of parasitism between virophage and giant virus, and between giant virus and host. Overexploitation by the giant virus would lead to extinction of the host (followed by virophage and giant virus, unless they expand their host range), while overexploitation by the virophage would lead to extinction of the giant virus (c.f., the Tragedy of the Commons). The latter would also reduce the possibility for horizontal transfer of the virophage (i.e., infection of new hosts) with potential long-term consequences for virophage fitness. The level of exploitation of a consumer depends on traits of the consumer (giant virus and virophage) and resource (host and giant virus) as well as densities and abiotic conditions. For long-term stability of the system, traits are predicted to evolve in a way that adaptations allow for sufficient exploitation without compromising the abundance of the host as a resource (9, 10). This balance may be difficult to evolve for the giant virus because different traits might determine the level of host exploitation and exploitation by the virophage, and negative associations between traits may constrain trait evolution. This has been shown for hyperparasitism (11–13), when hosts evolve in the presence of a virus/phage and predator (14, 15) or when hosts evolve in the presence of defensive and pathogenetic bacteria (16). For example, higher virulence of a pathogen and thus higher exploitation can evolve when virulence and the pathogen’s resistance against a competitor or a defensive microbe (or virophage) are positively correlated (17, 18). Lower virulence against the host was found when virulence is costly with respect to resistance against the defensive microbe (16, 17).

Here, we use experimental evolution to investigate processes that could contribute to the long-term persistence of host-virus-virophage systems by testing whether and how exploitation of giant viruses and virophages evolve over time. We established the marine bacterivore *Cafeteria burkhardae*, the giant virus CroV and the virophage mavirus (19–22) in chemostats (hereafter, we refer to CroV as ‘virus’ and to mavirus as ‘virophage’, (23)) and studied host population dynamics over 57 days either in the absence of virus and virophage, or with the virus, or with virus and virophage. To test for persistence of the three species and for evolutionary changes in the virus and virophage, we re-isolated virus and virophage from the chemostats after about ~280 host generations. We then compared virus and virophage replication and exploitation to their ancestors that were used to start the experiment.

Observations that the giant virus or virophage evolved greater exploitation would indicate that the system may become instable. Instability could lead to extinction or more pronounced oscillations in population dynamics and/or a longer time to reach equilibrium density, increasing the risk of extinction due to stochastic events. Long-term persistence would thus depend on specific environmental conditions that prevent overexploitation (e.g., spatial structure; 15). According to mathematical model analyses (7), persistence is more likely when virophage replication is slow, which can result from low intrinsic growth and/or high pathogenicity of virophages (= higher degree of giant virus exploitation). In addition, the giant virus is predicted to evolve a higher reproductive output in the absence of virophage and a lower reproductive output in the presence of virophage. Lower replication reduces the exploitation by the virophage and thus its replication and population size over time. The reproductive output of the giant virus is mainly determined by the intrinsic growth rate of the giant virus and/or virulence toward the host (= higher degree of host exploitation), as giant virus replication is associated with host cell lysis.

We found that host, virus, and virophage persisted for ~ 280 host generations, but that host population densities were not significantly different in the absence and presence of the virophage from ~ 140 generations onwards until the end of the experiment. We further observed that the virus evolved towards decreased replication while exploitation of the host did not change. The virophage had evolved increased replication and reduced inhibition of the virus, which results in decreased virus exploitation of the host.

## Materials and Methods

### Community dynamics experiment

Our experimental system consisted of the marine heterotrophic flagellate *Cafeteria burkhardae* strain E4-10P as host and the virus Cafeteria roenbergensis virus (CroV) and the virophage mavirus. The experiment was performed in chemostats (glass bottles with 400 mL f/2 enriched artificial seawater medium (50) supplemented with 0.025% (w/v) yeast extract and 0.3 g/mL of Chloramphenicol) with a continuous flow through of 120 ml medium per day (= 0.3 dilution rate per day) (23). Chemostats ran for 57 days. Replicated chemostats (n=3) were inoculated with the host and *Escherichia coli* bacteria (hereafter: control), with *E. coli* and the virus (hereafter: virus treatment), and with *E. coli*, virus and virophage (hereafter: virus-virophage treatment). All chemostats were inoculated with 7*10^4^ host cells/mL and virus and virophage were added at days 7 and 13 when host population dynamics had stabilized. Viruses and virophage were inoculated at a virus-to-host ratio of 0.1 in the virus and virus-virophage treatments. We used a Chloramphenicol resistant *E. coli* strain (BL21(DE3)pLysS) to keep the bacterial community constant. *E. coli* served as food for the host in our experiments.

We sampled chemostats (sample volume = 10 mL) every second day for host quantification. Host was quantified from live samples using a hemacytometer and light microscope (20x magnification). Virus samples were filtered (0.45 μm cellulose syringe filter) to remove host for later DNA extraction and quantification by digital droplet PCR (ddPCR), samples were kept at 4°C until measured. However, virus quantification failed because of significant losses during the filtration step.

For all ddPCR assays, DNA was extracted using a commercial kit (DNeasy 96 Blood & Tissue Kit, Qiagen, Hilden, Germany) and kept at 4°C until measured or measured right after sampling. We designed and selected primer and probes for virus amplification by standard PCR procedure (51). PCR parameters correspond to 1 cycle of 95°C (10 min), 40 cycle of 94°C (30 s), 40 cycle of 58°C (1 min), 1 cycle of 98°C (10 min) and hold temperature 12°C. All ddPCR results were analyzed using QUANTASOFT 1.7.4 The detailed methods and quality requirements for the data are described in the reference (52).

### Host-virus-virophage interactions

We amplified viruses from the virus treatment and virus-virophage treatment chemostats from the last sample day (day 57) of the experiment and used them to test for evolutionary changes. We used densities as proxies for the degree of virus and virophage exploitation and replication. To do this, we first separated day 57 viruses from host and bacteria and from each other by filtration through 0.45 μm and 0.2 μm filters. Viruses and virophages differ in size; CroV has a capsid size of 300 nm (53) and Mavirus of ~70 nm (21). We used a Spartan® cellulose filter (Whatman®) with a pore size of 0.2 μm to separate viruses and virophages from mixed samples. With a filter pore size of 0.2, CroV is mostly retained on the filter, while mavirus passes through the filter. We used reverse filtration to recover CroV from the filter (see below). To propagate and produce virus stocks of selected viruses, we added 50 μL of the medium used to recover viruses from the 0.2 μm filter to medium containing the ancestral host (strain E4-10P; 10^5^ cells/mL). After observing host lysis (confirmed by sampling and enumeration by microscopy), we collected virus samples (filtration through 0.45-μm filters) and quantified virus concentrations by ddPCR (51)). To produce selected virophages, we added 50 μL of the filtrate of the 0.2 μm filter to medium containing strain host (strain E4-10P; 10^5^ cells/mL) and ancestral virus. After observing host lysis, we collected virophage samples (filtration through 0.2 μm filter) and quantified virus concentrations by digital droplet PCR (ddPCR). Amplification of viruses and virophages also served as our evidence that viruses and virophages were present at the end of the chemostat experiment.

We tested whether viruses and virophages evolved during the chemostat experiment by comparing the replication and exploitation of ancestral and selected viruses and virophages. We infected the ancestral host (strain E4-10P) in 27 ml SW medium in tissue culture flasks with ancestral or selected viruses at a host-to-virus ratio of 0 (virus-free control) or 0.1; and with ancestral or selected virophages at a host-to-virus ratio of 0 (no virophage added control) or 10. All combinations were repeated three times starting with a host density of 10^4^ cells/mL. Host and virus samples were collected and quantified 1 day after infection. Host samples were preserved with glutaraldehyde (4% final concentration). Hosts were quantified by flow cytometry (FACSverse). For this purpose, samples were stained with SYBR-Green I DNA stain (Sigma-Aldrich) at a final concentration of 0.02%. Host enumeration was performed in well plates at a flow uptake of ~90 μl for 24 seconds with shaking at 1000 rpm. Lateral light scatter (SSC) and SYBR-green fluorescence (FITC) measurements were performed with logarithmic amplification using FlowJo software (BDBiosciences). Virus and virophage samples were immediately collected for DNA extraction and quantified by ddPCR (see above and (51)).

For reverse filtration, we attached a flexible tube (6 mm diameter) to the back of the filter and washed it with the medium used in the experiment to recover CroV. We measured the number of virophage DNA copies in the virus fraction (CroV) that were not removed from the virus fraction using this approach. On average, we detected ~10^3^ virophage DNA copies/mL, resulting in a virophage-host ratio of ~0.2 when considering the sample volumes added to achieve a virus-host ratio of 0.1 in the assays described above. In these assays, we detected amplification of virophages in 3 of 9 samples (2 replicate virus samples from chemostat 3, 1 from chemostat 9), but there were no differences in the densities of amplified viruses when comparing the densities of replicates with and without virophage amplification from the same chemostat. Thus, we conclude that the remaining virophages had no effect on the measurement of virus density 1 day after infection.

### Host-virus interaction

To test whether the host evolves resistance to the virus within a similar time frame as in the chemostat experiment, we performed an additional experiment running for 54 days. Host (strain E4-10P), ancestral virus, or ancestral virus and virophage were added to 30 mL of SW media in tissue culture flasks as 6 replicates. All communities started with a host density of 10^5^ cell/mL. Viruses were added at a virus/virophage-host ratio of 1. We transferred 8% of each community to a new flask with fresh medium every 4 days. After 54 days, we isolated clonal lines of the host from each flask. For this, we diluted the samples to 300 cell/mL and added 1 μL of the diluted sample to 200 μL medium in 96 well plates, we prepared one well plate per community. In addition, we prepared isolated clonal lines of the ancestor of the host (strain E4-10P) following the same protocol. After 6 days of growth, half of the isolated clonal lines per community and the ancestral host were infected with the ancestral virus to test for resistance to infection. The other half served as a virus-free control to account for the lack of growth independent of virus (no additional host, random extinction). At 8 days post-infection, host samples were fixated with Lugol’s solution (4% final concentration), and the presence or absence of hosts was later checked under a stereoscope. Resistance was estimated by comparing the percentage of wells that contained hosts between plates to which virus was added and virus-free controls.

### Virophage-host interaction: free and integrated virophage

We tested whether the number of virophages integrated into the genome and the presence of free virophages in the absence of the virus affected host growth using four different host strains: *C. burkhardae* strain E4-10P; strain RCC970-E3; two strains derived from the strain RCC970-E3. Strain RCC970-E3-8.8 and strain RCC970-E3-9.3.1 carry 2 or 3 copies of virophage integrated, respectively. Host growth was tested in the presence and absence of free virophages (virophage-host ratio of 8) in tissue culture flasks containing 30 ml SW medium and in 5 replicates (host: 2*10^3^ cell/mL). Samples were collected 6 days after infection, fixated (4% final concentration of glutaraldehyde), and host densities quantified by flow cytometry (see above).

### Data analysis

All data analyses were performed in Rstudio (54) and R (55) using the packages geepack (56) and lme4 (57). Differences between models were considered relevant when p<0.05 and significance was evaluated using model comparisons. All posthoc tests included corrections for multiple testing (Tuckey) using the package (multcomp) (58). To test for differences in host densities normalized to the average density of the controls without virus from the chemostat experiment, we used generalized linear models (GLM, family = quasipoisson) with treatment (host-virus, host-virus-virophage), period (day 1-34, day 35-57) and their interaction as explanatory variable (Fig. 1). We used GLM (family = poisson) to test for differences in virus densities to evaluate differences in virus replication and exploitation (Fig. 2A-C). For these tests, we used virus (ancestral, selected from the virus treatment, selected from the virus-virophage treatment) and virophage (absent, ancestral, selected) as well as their interaction as explanatory variables and chemostat as random effect to account for the relatedness of selected virus and virophage isolated from the same chemostat. To compare host densities in the presence of the virus, we used a GLM with virus (ancestral, selected from virus treatment, selected from virus-virophage treatment) as explanatory and chemostat as random variable (Fig. 2D). We used a GLM (family=poisson) to test for differences in virophage density with virophage (ancestral, selected), virus (ancestral, selected from the virus treatment, selected from the virus-virophage treatment) and their interaction as explanatory and chemostat as random variable (Fig. 3). We compared host densities in the presence of virus and presence and absence of virophage (Fig. 4) using a GLM (family=poisson) with virus (ancestral, selected from the virus treatment, selected from the virus-virophage treatment), virophage (ancestral, selected) and their interaction as explanatory variable. To test for differences in host resistance, we compared the percentage of wells with host growth in the presence or absent of ancestral virus using a GLM (family=poisson) with virus (presence, absence) and host line (ancestral host, host isolated from communities without virus, host isolated from communities with virus, host isolated from communities with virus and virophage) as explanatory variable. We used well plates as random effect. For the virophage-host interaction experiment (free and integrated virophage, Fig. S3), we compared the average host growth rate with host lineage (host strains) and virophage (presence, absence) as the explanatory variables using generalized linear models (GLM, family = gaussian).

**Figure 1.**
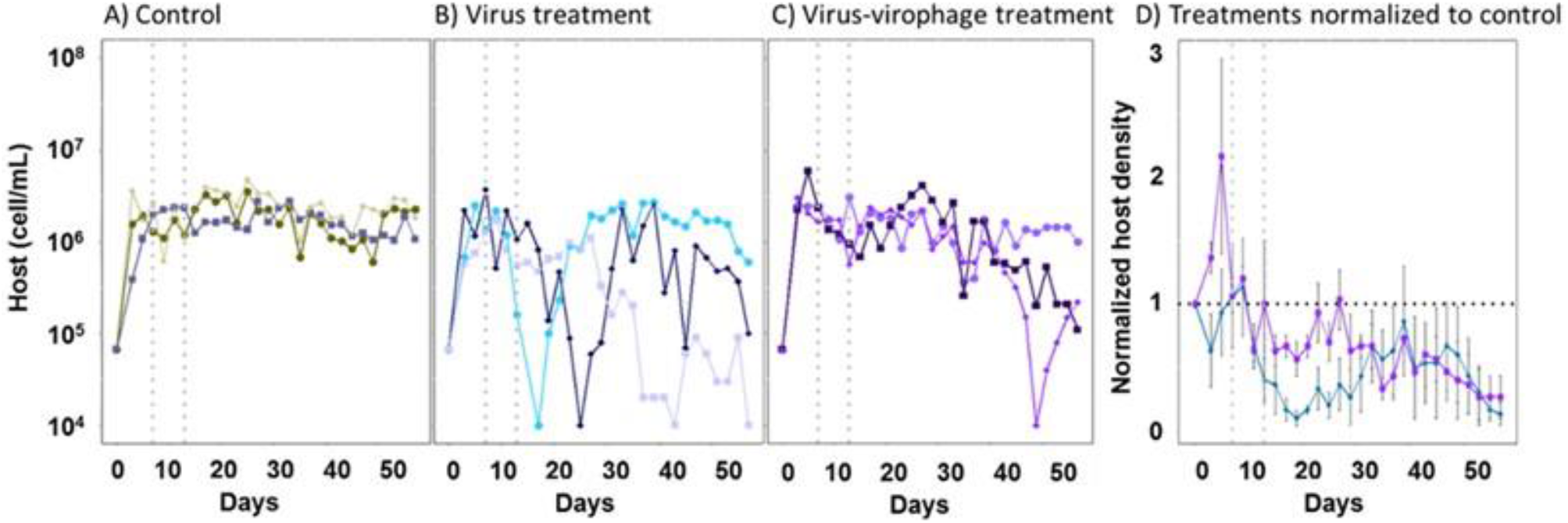
Persistence of tripartite system. Host population density (cell/mL) dynamics in chemostats A) without virus and virophage, B) with virus and without virophage, C) with virus and virophage (each line shows a single replicate, n=3). Grey vertical lines in A-C indicate addition of virus and virophage. D) Host population density in the presence of the virus (blue) as well as virus and virophage (purple) normalized with respect to the host density in the absence of viruses (black horizontal line; mean ± s.d., n=3).

**Figure 2.**
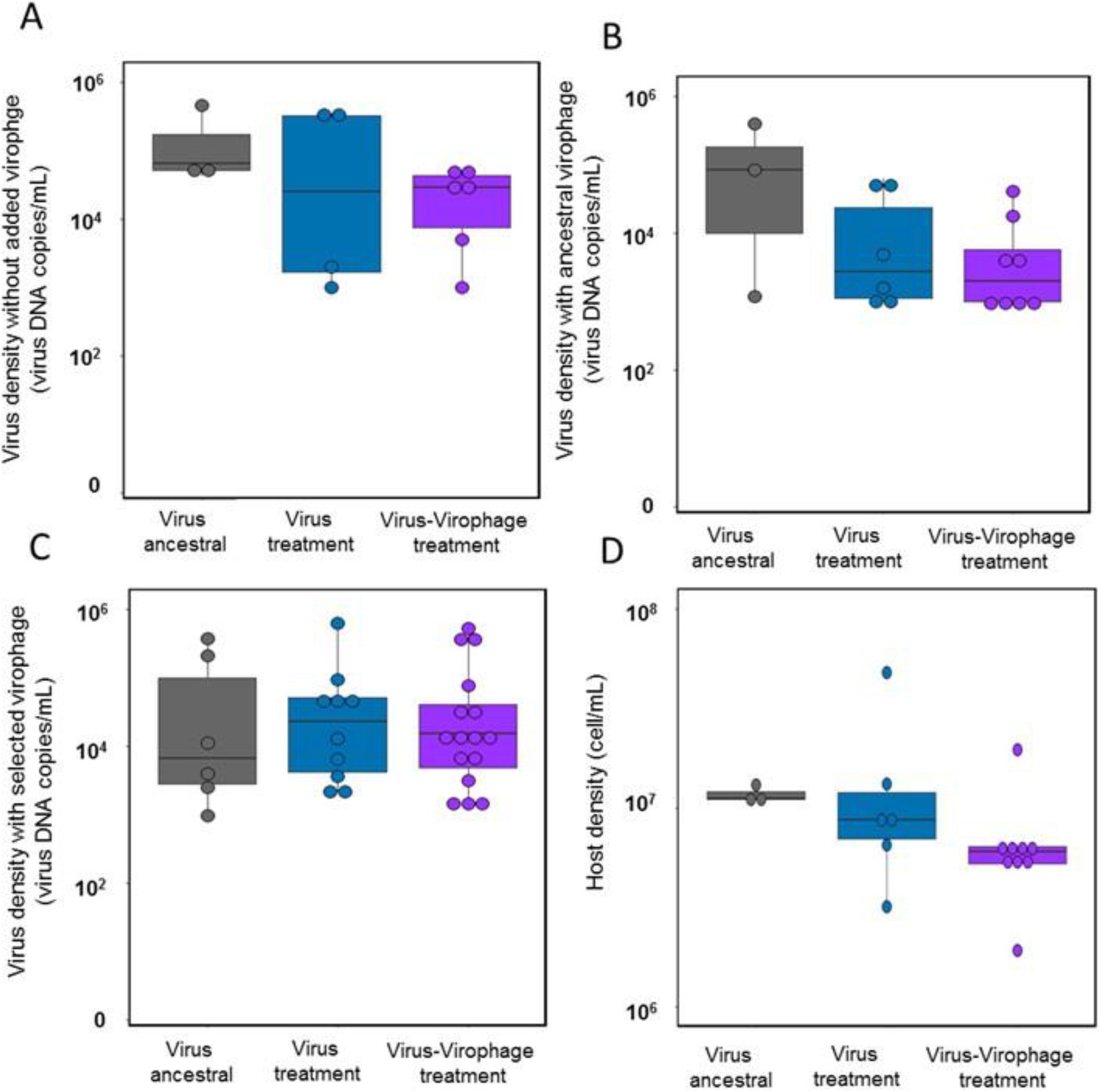
Changes in virus replication and host exploitation. Density of ancestral and selected virus lines (as viral DNA copy numbers/mL, median). Virus densities differed between virus lines and the presence of the virophage being overall lower for selected viruses: (A), virus density without virophage, (B) virus density with virophage ancestral, (C) virus density with virophage selected lines, and (D), host exploitation as host density with virus ancestral or selected lines. Grey (ancestral), blue (selected virus from the chemostat with virus), purple (selected virus from the chemostat with virus and virophage). There were 3 replicates per virus type (either ancestral or selected) being: 1 ancestral virus, 2 selected virus coming from the virus-treatment, 3 selected virus coming from the virus-treatment and 3 selected virophage coming from the virus-virophage treatments.

**Figure 3.**
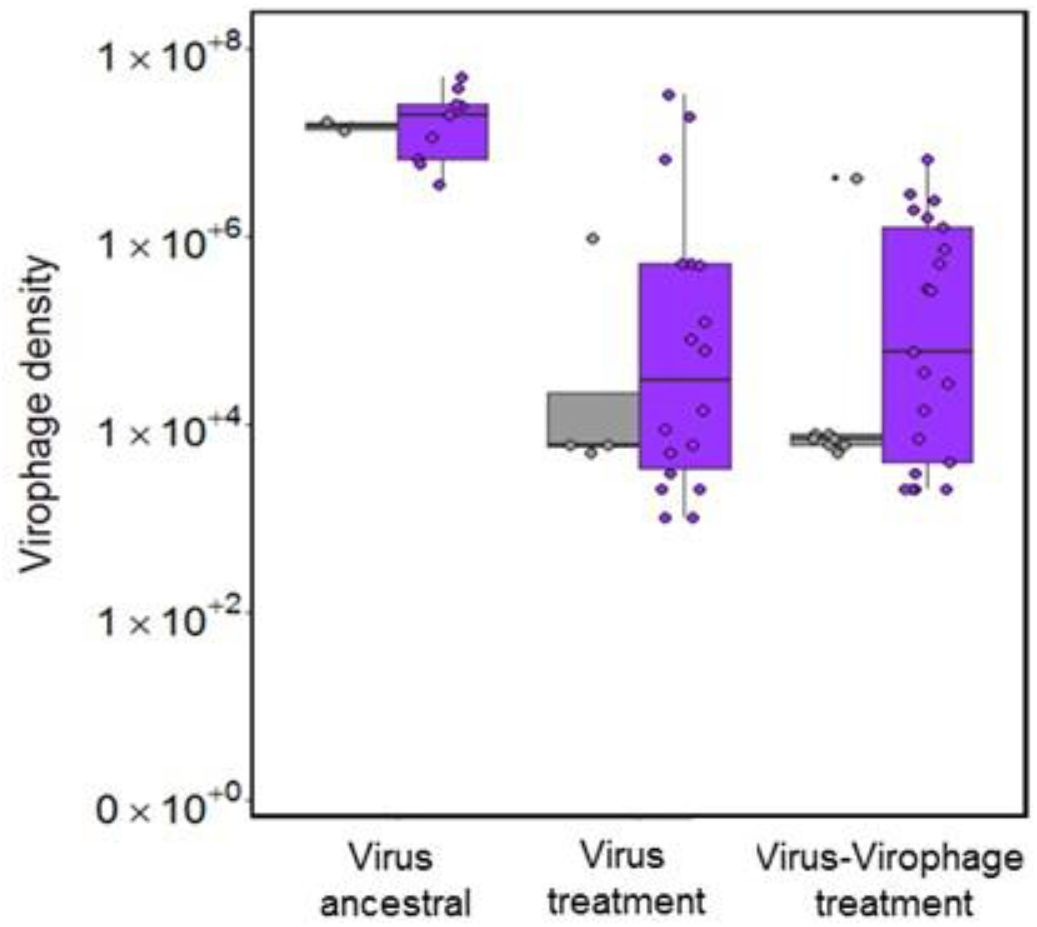
Changes in virophage replication. Virophage density (as virophage DNA copy numbers/mL, median) of ancestral (grey) and selected virophage lines (purple) in the presence of ancestral virus 1 day post infection.

**Figure 4.**
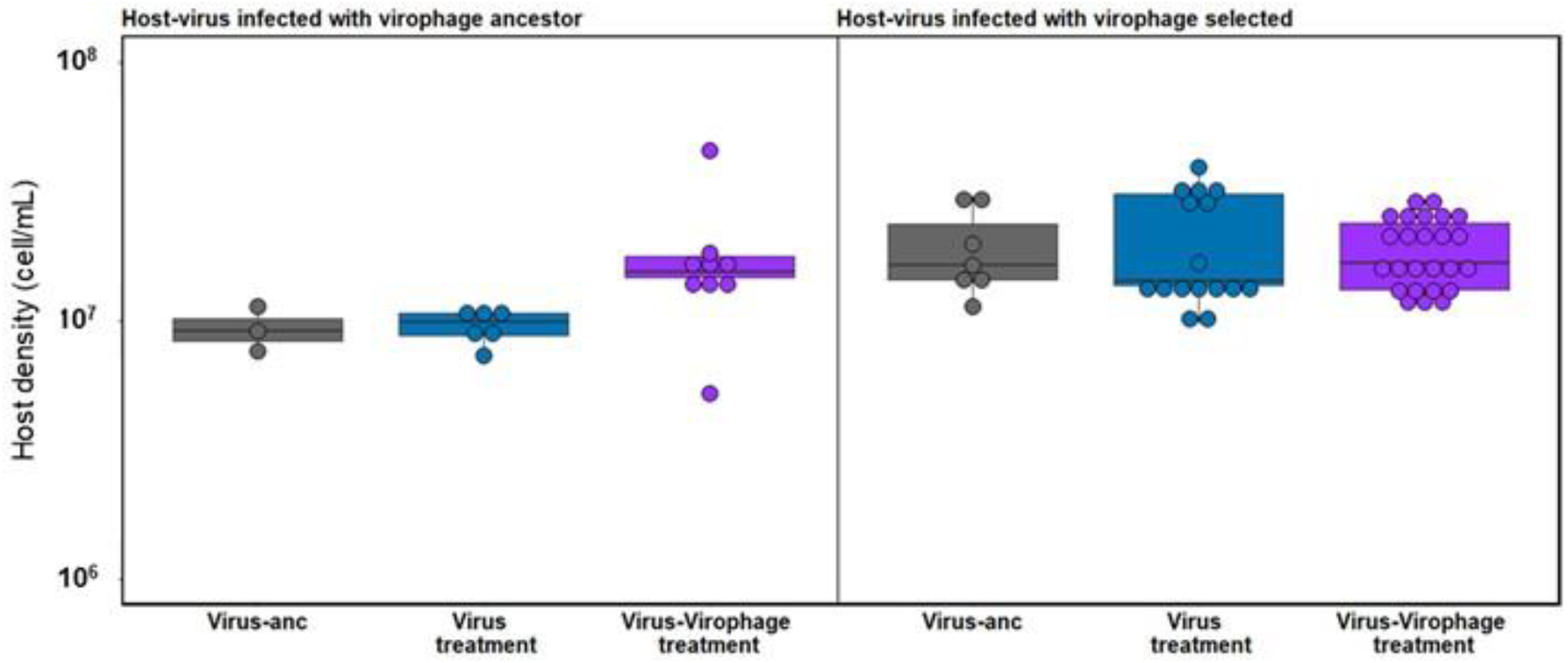
Effect of virophage on host exploitation by the virus. Host density (as cell/mL, median; ancestral host) when grown for 1 day in the presence of virus (ancestral: grey; selected from virus treatment: blue; virus-virophage treatment: purple) and virophage (left panel: ancestral virophage; right panel: selected virophage). Host densities were significantly higher in the presence of virus from the virus-virophage treatment with ancestral virophage (left panel) and in the presence of the selected virophage (right panel; for statistics see main text).

## Results

We investigated the long-term persistence of host-virus-virophage systems to test whether and how replication and exploitation of giant viruses and virophages evolve. In this system, virus infection of the host nearly always leads to cell lysis. During a virophage-only infection of the host, the virophage may integrate into the nuclear genome and reproduce as part of the host genome (25). When virus and virophage synchronously coinfect a host cell, virophage production blocks virus reproduction and only virophage is produced, which mitigates the effect of the giant virus on the host population during subsequent infections. When the virus infects a host with integrated virophage in their genome, both virus and virophage are produced, with no apparent decrease in virus replication.

We grew hosts over 57 days in replicated chemostats (continuous culture systems, n=3 per community) in the absence of virus and virophage (control), with the virus (virus treatment), and with virus and virophage (virus-virophage treatment). Following standard experimental evolution protocols with microbes, we re-isolated virus and virophage populations from the end of the experiment and assayed virus and virophage traits after ~280 host generations (number of generations under optimal replication conditions of the host) and of the ancestor. The flagellate host strain used to start the experiments carried endogenous virophages (4), but these did not amplify under the experimental conditions of this study (Supplementary information Fig. S1) and we thus consider the control and the virus treatment to lack free virophages. We confirmed the absence of free virophage from the control and virus treatment at day 57.

### Host dynamics and persistence

We followed host population sizes over 57 days and confirmed the presence of virus and virophage at day 57 through amplification from subsamples (see below: isolation of selected virus and virophage). We found that the host-virus-virophage could persist over the course of the experiment without driving each other to extinction but we observed significant differences in the temporal dynamics of the host population. Host populations decreased rapidly after inoculation of the virus in the virus and virus-virophage treatments (Fig. 1B,C) but less so when the virophage was present (Fig. 1C). Host populations recovered in two out of three replicates of the virus-treatment before declining again, resembling consumer resource cycles (26). In the communities with the virus and virophage, host population sizes remained relatively stable and declined only towards the end of the experiment (Fig. 1C) while host population remained at ≈10^6^ cells mL^−1^ throughout the experiment in the absence of virus and virophage (Fig. 1A, controls). As we observed differences in the temporal trends in the virus and virus-virophage treatments and to uncover the potential role of the virophage for host population dynamics, we normalized host densities from the virus and virus-virophage treatments to the average host density of the replicates from the control treatment. According to the predicted beneficial effect of the virophage for the host population, the normalized host densities were significantly lower in the virus treatment compared to the virus-virophage treatment but only during the first half of the experiment. This difference in normalized host densities was lost as host densities became more similar across treatments starting around mid-time point of the experiment (~140 host generations, 35 days; Generalized linear model (GLM): treatment: df=2, X^2^=5,72, p=0.0005; period: df=2, X^2^=7.48,p=4.3*10^−5^; interaction treatment and period (days 1-34, 35-57): df=1, X^2^=2.75, p=0.007).

### Virus and virophage isolation

We isolated virus and virophage from the chemostats at day 57 of the experiment by adding a subsample from each chemostat to ancestral host populations that were used to inoculate the chemostats. The amplification took approx. 21 generations of virus replication and approx. 2 generations of virophage replication after the isolation from the chemostats (see material and methods for details). In this way we amplified virus populations from all communities that were inoculated with virus, and virophage populations from those where we added virophage at the beginning of the experiment. We refer to *selected* virus and virophage lines for the amplified viruses and virophages, and *ancestral* virus and virophage lines to the virus and virophage that were used to inoculate the chemostats. Following standard experimental evolution protocols with microbes, we used these viruses to measure levels of exploitation and replication under standardized conditions. This allowed us to identify heritable phenotypic changes in the virus populations (15, 27). Specifically, we used host densities and virus/virophage DNA copy numbers one day post infection as proxies for levels of exploitation and replication. The initial ratios of virus and host, the virus and virophage as well as densities of hosts were the same across assays, and we exclude density dependent effects in these assays. We used the ancestral host population for all experiments as there was no evidence that hosts evolved resistance to the virus (see below).

### Virus evolution

We compared virus densities of ancestral and selected virus (virus and virus-virophage treatment) in the presence or absence of virophage (ancestral or selected) one day post infection. We found that ancestral and selected virus densities differed significantly and depended on whether ancestral or selected virophage was present (Generalized linear mixed model (GLMER): virus χ2= 366324, df=6, p<2.2*10^−16^; virophage χ2= 7760494, df=6, p<2.2*10^−16^; virophage χ2= 7760494, df=6, p<2.2*10^−16^; interaction virus and virophage χ2= 366321, df=4, p<2.2*10^−16^, Fig. 2). Specifically, we found that virus density was lower for selected virus from the virus-virophage treatment compared to the ancestral virus (Posthoc test: p<0.0001; all other comparisons non-significant, Fig. 2A). Virus density was significantly lower in presence of the ancestral virophage, compared to the selected virophage (Posthoc test: ancestor and virophage absent: p<2.2*10^−16^; selected and virophage absent: p<2.2*10^−16^; ancestral and selected virophage: p<2.2*10^−16^, Fig. 2B,C).

To further assess host exploitation by the virus, we compared host densities one day post infection in the presence of ancestral and selected virus (virus and virus-virophage treatment). Host densities were not significantly different depending on the virus (GLMER: χ2= 3.94, df=2, p=0.139, Fig. 2D), even if there was a trend towards lower densities for selected viruses from the chemostats with virophage present (Posthoc test: selected virus-virophage and selected virus treatment: p=0.069).

### Virophage evolution

Ancestral virophage density was significantly higher in the presence of the ancestral virus compared to the selected virus, and virophage density was higher for selected virophage than for ancestral virophages when tested with ancestral and selected virus (Fig. 3, LME: interaction virus and virophage line, χ^2^= 10949652, df=2, p < 2.2 *10^−16^; virus line, χ^2^= 10949655, df=4, p < 2.2*10^−16^; virophage line, χ^2^=42586791, df=3, p < 2.2*10^−16^). To further assess how host exploitation by the virus is affected by the virophage, we compared host densities in the presence of ancestral and selected virus (virus and virus-virophage treatment) and presence of ancestral or selected virophage. Host density was significantly affected by the combination of virus and the virophage lines (GLMER: interaction virus and virophage line, χ^2^= 18317680, df= 2, p < 2.2 *10^−16^; virus line, χ^2^= 18317684, df=4, p < 2.2*10^−16^; virophage line, χ^2^= 42042860, df=3, p < 2.2*10^−16^, Fig. 4). Specifically, host densities with selected virus from the virus-virophage treatment had significantly higher densities compared to the other two virus lines (Posthoc test: ancestral virus vs selected in presence of virus and virophage: p<0.0001; selected in presence of virus vs selected in presence of virus and virophage: p<0.0001; ancestral virus vs selected in presence of virus: p=0.978, Fig. 4) and host densities were highest in the presence of the selected virophage from the virus-virophage treatment (Posthoc test: ancestral virus vs selected virus from the virus and virus-virophage treatment: p < 2.2*10^−16^; selected virus from the virus-treatment vs selected virus from the virus-virophage treatment: p < 2.2*10^−16^; ancestral virus vs selected virus from the virus-treatment: p < 2.2 *10^−16^, Fig. 4).

### Host evolution

We tested for the potential of the host evolving resistance against the virus in a separate experiment. We isolated hosts after 54 days from semi-continuous cultures where hosts grew in the absence of the virus, the presence of the virus, and the virus and virophage. We then compared growth of ancestral and isolated hosts (54 days) in the absence and presence of ancestral virus. Growth of hosts was significantly reduced in the presence of the virus and independent of the host line (Supplementary information Fig. S2, LME, interaction virus presence and host line, df=3, χ^2^=15.39, p=0.001; host line, df=6, χ^2^=19.22, p=0.004; virus presence, df=4, χ^2^=15.43, p=0.004). Hence, we did not observe any host resistance to the virus in these experiments.

### Host-virophage interaction

We assessed if the number of virophages integrated in the genome and the presence of free virophages affected host growth in the absence of the virus by using host strains with different numbers of virophage copies integrated in their genome. We found no effect of integrated or free virophage on host strain growth rate (Supplementary information Fig. S3, LME, interaction host line and virophage type, χ^2^=-0.73, df= 3, p=0.975)

## Discussion

Long-term persistence of consumer and resource (i.e., host-parasite, predator-prey) is only possible if the consumer does not exploit the resource beyond the point where sustainable exploitation is no longer possible. This problem appears to be prominent in systems where one species exploits a resource while being exploited by another species, since adaptation must favor traits that balance exploitation with avoidance of being exploited. Because trait evolution can be constrained for energetic or genetic reasons (28), selection often cannot optimize all traits simultaneously, with potential consequences for species persistence. We studied here persistence and tracked changes in exploitation in a host-virus-virophage system to test whether exploitation evolves to promote persistence (7). We did this by following host population dynamics in long-term coculture experiments and isolating viruses from the last experimental day and found that the three species persisted over the course of the experiment. We did not follow virus and virophage densities over time and we do thus have no information on virus and virophage replication. It is theoretically possible, but highly unlikely given the continuous dilution of chemostat cultures (dilution rate = 0.3 d^−1^), that virus and virophage amplified from the end of the experiment are residual ancestral virus and virophage or were produced at the beginning of the experiment rather than virus and virophage produced throughout the experiment. This conclusion is supported by the observation that virus and virophage replication and exploitation differs when comparing ancestral and selected populations in standardized assays.

We used the isolated viruses from day 57 to evaluate changes in exploitation and replication by comparing densities of host, virus and/or virophage one day post infection when combining host, virus and virophage from the start of the experiment and after ~ 280 host generations coculturing. We found that virus evolved lower replication while host exploitation did not change. The virophages evolved increased replication and lower exploitation of the virus. Finally, we observed that the selected virophage led to decreased host exploitation by the virus when tested with ancestral and selected virus. Lower virus replication as well as lower virus inhibition by the virophage can promote persistence as these reduce the impact of the virus on the host population and the possibility of virophage extinction due to oscillatory dynamics linked to high inhibition of the virus (7).

The host-virophage interaction became less beneficial for the host over time in our experiment. At least two non-exclusive mechanisms could explain this observation. First, the evolutionary changes in exploitation and replication of the giant virus and virophage could lead to changes in the observed differences in the temporal dynamics of the host population over time. Reduced exploitation can stabilize consumer resource interactions (29) and evolutionary changes in the level of exploitation can stabilize consumer-resource dynamics (30). Second, the most common mode of infection by the virophage (reactivation vs. coinfection) could affect the role of the virophage for the host population as well as the selection on the virophage. An increasing frequency of hosts carrying virophage over time in the presence of the virus might favor selection that maximizes the number of coinfections. This is related to selection on bacteriophages and other parasites (31–33) or mutualists (34, 35) with horizontal or vertical transmission where the relative opportunity for horizontal versus vertical transmission has been shown to affect the evolution of symbiont effects on host fitness (e.g. (36–41)). Generally, horizontal transmission favors increased costs of infection for the host and more virulent phages or less beneficial mutualists, which leads to an increase in the rate of infection transfers. Virophage transmission is only vertical in the absence of giant virus infection in the present system, which is expected to be the major mode of transmission in wild host populations, as evidenced by the widespread occurrence of integrated virophages (4). As the giant virus was present at the end of the experiment, we expect that transmission was predominantly horizontal under the experimental conditions. Further studies are needed to investigate the role of infection mode frequency and its consequences for selection and evolution.

In our experimental community, the host did not evolve resistance to the giant virus independent of whether the virophage was present or not. Host-associated microbes that mitigate the effects of pathogens on the host are common in nature and have been shown to shape evolutionary dynamics, reducing selection for resistance in the host (16, 42, 43). For example, the evolution of bacteriophages and satellite viruses showed that the coevolution of helper bacteriophages and satellite viruses reduces selection for bacterial resistance to the bacteriophage (44–46). The contribution of coevolution of virus and virophage to constraining resistance evolution is, however, unclear in our experiment, as we also did not observe host resistance evolution in the absence of virophage, which may be due to the relatively short duration of the experiment. We found some evidence of virus-virophage coevolution; virus replication decreased in the presence of ancestral and selected virophages for virus from the virus-virophage treatment, and host exploitation was lowest when virus and virophage from the virus-virophage treatment were combined. However, more detailed experiments and tests comparing the replication and exploitation of host, virus, and virophage when paired from different time points in the experiment are needed. They could for example provide insights into the pattern of coevolution (27, 47, 48).

The experiments were started with non-clonal populations of giant viruses and virophages and the changes we observed could thus be the result of selection on standing genetic variation and/or on de-novo mutations occurring in the chemostats. This is, however, not important for the interpretation of the results discussed here, as in both cases, we can assume that the frequency of the genotypes was low (or absent) at the start of the experiment. Similar to other experimental host-pathogen systems (27, 31, 49), the present system had a high potential for coevolution of traits determining species interactions. In particular, the small genome size of the viruses, the large population sizes (estimated harmonic mean population size from an independent experiment for viruses 2*10^4^ cells/mL and virophages 3*10^6^ particles/mL; del Arco *unpublished data*), and the strong selection pressure (27) make de novo mutations likely to occur in this experiment.

Identifying and understanding the processes that allow interacting species with strong interdependence and mutual exploitation to persist is an ongoing challenge, and yet there are many such systems that persist over time. One evolutionary strategy that has been identified to facilitate persistence of such systems is reduced exploitation and reproduction. We showed in our experiments, that virus and virophage evolved reduced degrees of exploitation and replication confirming general theory on coexistence in tripartite systems and proving experimental evidence for processes that could facilitate persistence of protist-giant virus-virophage systems.

## Author contribution

AdA, LB, MF conceived the study, AdA, LB designed of the study; AdA carried out the experiment, AdA, LB analysed the data; AdA, LB wrote the manuscript, all authors edited the manuscript.

## Competing Interest Statement

We have no competing interests.

## Acknowledgments

This work was funded by Gordon and Betty Moore Foundation to MGF and LB (Grant #5734). We thank Ruben Herrmann for help with the establishment of the reverse filtration protocol and the Fischer Lab for helpful comments on the first draft of the manuscript.

## Notes

### Competing Interest Statement

The authors have declared no competing interest.

## References

1. S. Duponchel, M. G. Fischer, Viva lavidaviruses! Five features of virophages that parasitize giant DNA viruses. PLOS Pathogens 15, e1007592 (2019).

2. D. Paez-Espino, et al., Diversity, evolution, and classification of virophages uncovered through global metagenomics. Microbiome 7, 157 (2019).

3. M. Berjón-Otero, A. Koslová, M. G. Fischer, The dual lifestyle of genome-integrating virophages in protists. Ann N Y Acad Sci 1447, 97–109 (2019).

4. T. Hackl, S. Duponchel, K. Barenhoff, A. Weinmann, M. G. Fischer, Virophages and retrotransposons colonize the genomes of a heterotrophic flagellate. eLife 10, e72674 (2021).

5. M. G. Fischer, T. Hackl, Host genome integration and giant virus-induced reactivation of the virophage mavirus. Nature 540, 288–291 (2016).

6. E. V. Koonin, M. Krupovic, A parasite’s parasite saves host’s neighbours. Nature 540, 204–205 (2016).

7. D. Wodarz, Evolutionary dynamics of giant viruses and their virophages. Ecol Evol 3, 2103–2115 (2013).

8. R. D. Holt, M. E. Hochberg, The Coexistence of Competing Parasites. Part II—Hyperparasitism and Food Chain Dynamics. Journal of Theoretical Biology 193, 485–495 (1998).

9. S. Heilmann, K. Sneppen, S. Krishna, Sustainability of Virulence in a Phage-Bacterial Ecosystem. J Virol 84, 3016–3022 (2010).

10. A. Y. Morozov, C. Robin, A. Franc, A simple model for the dynamics of a host-parasite-hyperparasite interaction. J Theor Biol 249, 246–253 (2007).

11. H. S. Addy, A. Askora, T. Kawasaki, M. Fujie, T. Yamada, Loss of Virulence of the Phytopathogen Ralstonia solanacearum Through Infection by φRSM Filamentous Phages. Phytopathology® 102, 469–477 (2012).

12. R. M. May, M. P. Hassell, The Dynamics of Multiparasitoid-Host Interactions. The American Naturalist 117, 234–261 (1981).

13. S. R. Parratt, A.-L. Laine, The role of hyperparasitism in microbial pathogen ecology and evolution. ISME J 10, 1815–1822 (2016).

14. J. Frickel, L. Theodosiou, L. Becks, Rapid evolution of hosts begets species diversity at the cost of intraspecific diversity. Proceedings of the National Academy of Sciences 114, 11193–11198 (2017).

15. P. Gómez, A. Buckling, Real-time microbial adaptive diversification in soil. Ecology Letters 16, 650–655 (2013).

16. S. A. Ford, D. Kao, D. Williams, K. C. King, Microbe-mediated host defence drives the evolution of reduced pathogen virulence. Nat Commun 7, 13430 (2016).

17. M. León, R. Bastías, Virulence reduction in bacteriophage resistant bacteria. Front. Microbiol. 6 (2015).

18. J. Santander, J. Robeson, Phage-resistance of Salmonella enterica serovar Enteritidis and pathogenesis in Caenorhabditis elegans is mediated by the lipopolysaccharide. Electron. J. Biotechnol. 10, 0–0 (2007).

19. A. Schoenle, Global comparison of bicosoecid Cafeteria-like flagellates from the deep ocean and surface waters, with reorganization of the family Cafeteriaceae. European Journal of Protistology, 21 (2020).

20. T. Fenchel, D. J. Patterson, Cafeteria roenbergensis nov. gen., nov. sp., a heterotrophic microflagellate from marine plankton (1988) (December 2, 2021).

21. M. G. Fischer, C. A. Suttle, A Virophage at the Origin of Large DNA Transposons. Science 332, 231–234 (2011).

22. M. G. Fischer, M. J. Allen, W. H. Wilson, C. A. Suttle, Giant virus with a remarkable complement of genes infects marine zooplankton. Proceedings of the National Academy of Sciences of the United States of America 107, 19508–13 (2010).

23. A. del Arco, N. Woltermann, L. Becks, Building up chemostats for experimental eco-evolutionary studies (2020) https://doi.org/10.17504/protocols.io.tkxekxn (November 16, 2020).

24. M. A. Nowak, R. M. May, Virus dynamics: Mathematical principles of immunology and virology (Oxford University Press, 2001).

25. M. G. Fischer, T. Hackl, Host Genome Integration and Reactivation of the Virophage Mavirus In the Marine Protozoan Cafeteria roenbergensis. Nature 540, 288–291 (2016).

26. E. Pachepsky, R. M. Nisbet, W. W. Murdoch, Between Discrete and Continuous: Consumer–Resource Dynamics with Synchronized Reproduction. Ecology 89, 280–288 (2008).

27. J. Frickel, M. Sieber, L. Becks, Eco-evolutionary dynamics in a coevolving host–virus system. Ecology Letters 19, 450–459 (2016).

28. G. de Jong, A. J. van Noordwijk, Acquisition and Allocation of Resources: Genetic (CO) Variances, Selection, and Life Histories. The American Naturalist 139, 749–770 (1992).

29. M. L. Rosenzweig, R. H. MacArthur, Graphical Representation and Stability Conditions of Predator-Prey Interactions. The American Naturalist 97, 209–223 (1963).

30. T. Hiltunen, et al., Dual-stressor selection alters eco-evolutionary dynamics in experimental communities. Nat Ecol Evol 2, 1974–1981 (2018).

31. J. W. Shapiro, E. S. C. P. Williams, P. E. Turner, Evolution of parasitism and mutualism between filamentous phage M13 and Escherichia coli. PeerJ 4 (2016).

32. S. Fellous, L. Salvaudon, How can your parasites become your allies? Trends Parasitol 25, 62–66 (2009).

33. M. Lipsitch, S. Siller, M. A. Nowak, The evolution of virulence in pathogens with vertical and horizontal transmission. Evolution 50, 1729–1741 (1996).

34. J. W. Shapiro, P. E. Turner, The impact of transmission mode on the evolution of benefits provided by microbial symbionts. Ecology and Evolution 4, 3350–3361 (2014).

35. M. J. Roossinck, The good viruses: viral mutualistic symbioses. Nature Reviews Microbiology 9, 99–108 (2011).

36. R. M. May, R. M. Anderson, Epidemiology and genetics in the coevolution of parasites and hosts. Proc. R. Soc. Lond. B. 219, 281–313 (1983).

37. P. W. Ewald, Transmission modes and evolution of the parasitism-mutualism continuum. Ann N Y Acad Sci 503, 295–306 (1987).

38. J. J. Bull, I. J. Molineux, W. R. Rice, Selection of benevolence in a host-parasite system. Evolution 45, 875–882 (1991).

39. J. J. Bull, VIRULENCE. Evolution 48, 1423–1437 (1994).

40. D. Ebert, Virulence and local adaptation of a horizontally transmitted parasite. Science 265, 1084–1086 (1994).

41. T. Day, Parasite transmission modes and the evolution of virulence. Evolution 55, 2389–2400 (2001).

42. K. M. Oliver, A. H. Smith, J. A. Russell, Defensive symbiosis in the real world – advancing ecological studies of heritable, protective bacteria in aphids and beyond. Functional Ecology 28, 341–355 (2014).

43. S. Polin, J.-C. Simon, Y. Outreman, An ecological cost associated with protective symbionts of aphids. Ecol Evol 4, 826–830 (2014).

44. G. E. Christie, T. Dokland, Pirates of the Caudovirales. Virology 434, 210–221 (2012).

45. M. Krupovic, J. H. Kuhn, M. G. Fischer, A classification system for virophages and satellite viruses. Archives of Virology 161, 233–247 (2016).

46. B. Frígols, et al., Virus Satellites Drive Viral Evolution and Ecology. PLOS Genetics 11, e1005609 (2015).

47. S. Gaba, D. Ebert, Time-shift experiments as a tool to study antagonistic coevolution. Trends in Ecology and Evolution 24, 226–232 (2009).

48. J. Frickel, L. Theodosiou, L. Becks, Rapid evolution of hosts begets species diversity at the cost of intraspecific diversity. PNAS 114, 11193–11198 (2017).

49. B. Koskella, M. A. Brockhurst, Bacteria-phage coevolution as a driver of ecological and evolutionary processes in microbial communities. FEMS Microbiology Reviews 38, 916–931 (2014).

50. R. R. Guillard, J. H. Ryther, Studies of marine planktonic diatoms. I. Cyclotella nana Hustedt, and Detonula confervacea (cleve) Gran. Can J Microbiol 8, 229–239 (1962).

51. A. del Arco, M. Fischer, L. Becks, Simultaneous Giant Virus and Virophage Quantification Using Droplet Digital PCR. Viruses 14, 1056 (2022).

52. H. Koch, J. Frickel, M. Valiadi, L. Becks, Why rapid, adaptive evolution matters for community dynamics. Frontiers in Ecology and Evolution 2, 1–10 (2014).

53. C. Xiao, et al., Cryo-EM reconstruction of the Cafeteria roenbergensis virus capsid suggests novel assembly pathway for giant viruses. Sci Rep 7, 5484 (2017).

54. RStudio Team, RStudio: Integrated Development for R. RStudio, PBC, Boston, MA URL http://www.rstudio.com/. (2020).

55. R Core Team, R: A language and environment for statistical computing. R Foundation for Statistical Computing, Vienna, Austria. URL https://www.R-project.org/. (2020).

56. S. Højsgaard, U. Halekoh, J. Yan, C. Ekstrøm, geepack: Generalized Estimating Equation Package (2020) (January 11, 2021).

57. J. Pinheiro, D. Bates, S. DebRoy, D. Sarkar, R. C. Team, nlme: Linear and Nonlinear Mixed Effects Models. R package version 3.1-131. 69, 18637 (2017).

58. T. Hothorn, F. Bretz, P. Westfall, Simultaneous Inference in General Parametric Models. Biometrical Journal 50, 346–363 (2008).

