## Supplementary material for "Evolution of virus and virophage facilitates persistence in a tripartite microbial system": bioRxiv_delArco_si.pdf

### Supplementary Information

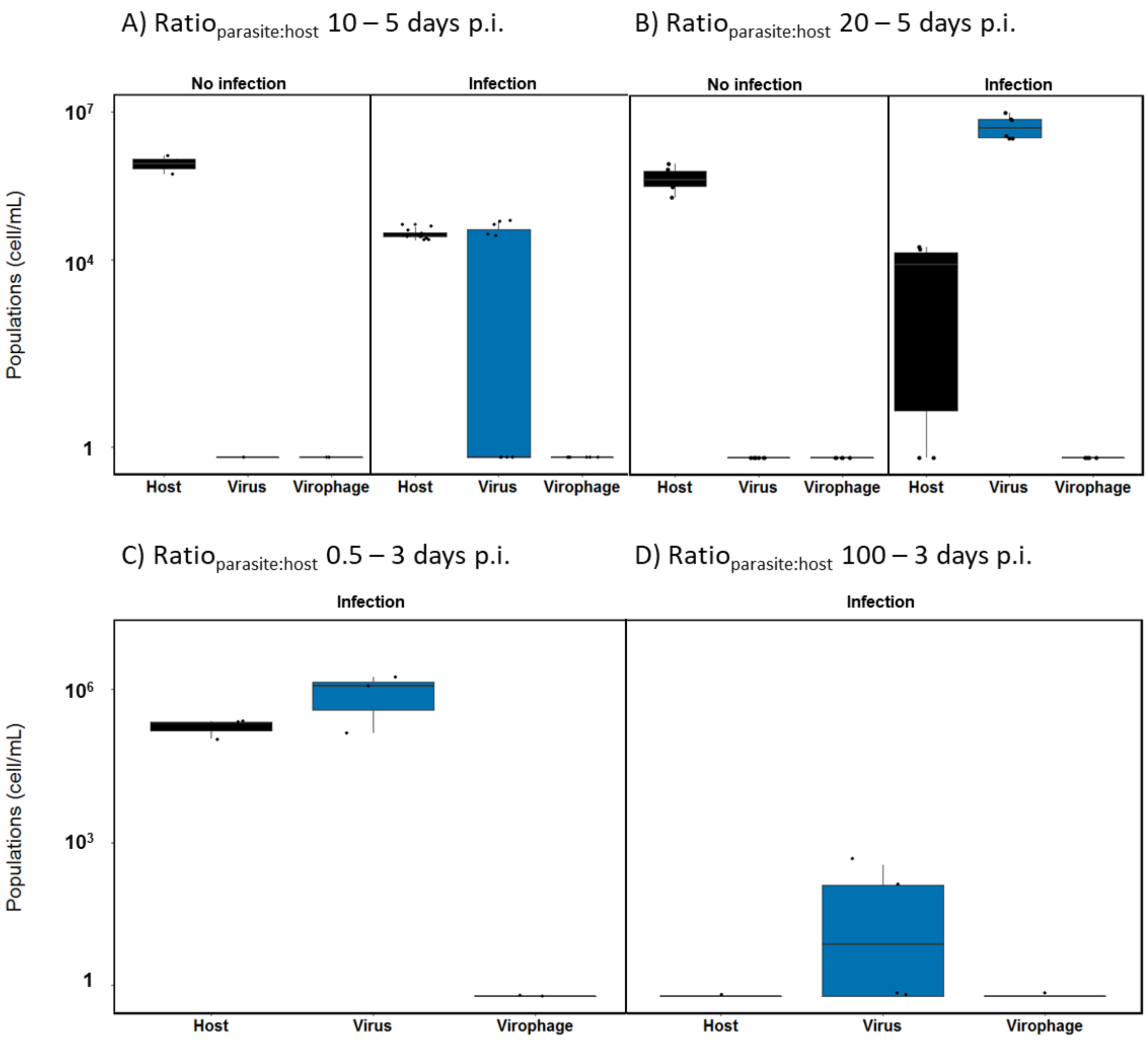


**Fig. S1.** Assessment of virophage reactivation from host ancestral genome under infection. Virophage reactivation was not detected. Host, virus and virophage population density (cell/mL) under a range of infection conditions varying in virophage:virus ratio ($Ratio_{parasite:host}$) and duration (A) $Ratio_{parasite:host}$ 10 – 5 days post infection (p.i., n=4 shown by points), (B) $Ratio_{parasite:host}$ 20 – 5 days p.i. (n=6, shown by points), (C) $Ratio_{parasite:host}$ 0.5 – 3 days p.i. (n=3, shown by points) and (D) $Ratio_{parasite:host}$ 100 – 3 days p.i. (n=3, shown by points). Grey (host), blue (virus), purple (virophage).

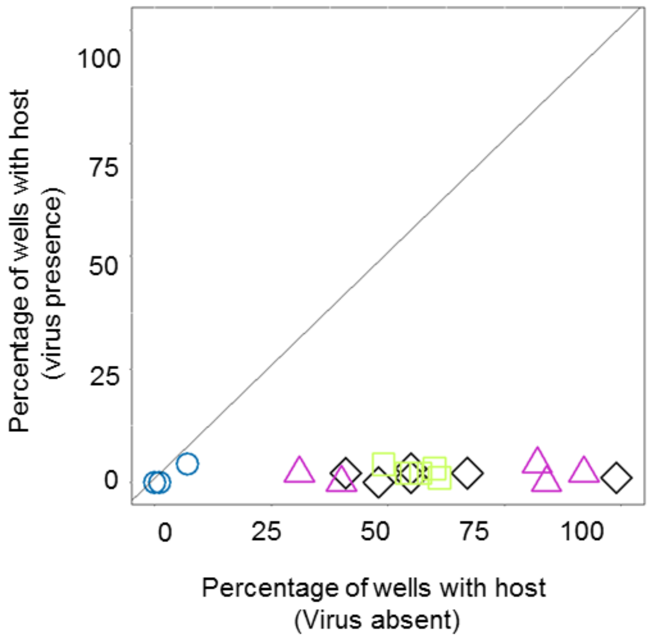


**Fig. S2.** Comparison of host growth as percentage of wells where host clones grew in the absence (x-axis) and presence of the ancestral virus (y-axis). Symbols represent host lines: ancestral host (black diamonds), hosts from chemostats without virus and virophage (green squares), host from chemostat with virus (blue circles), and host from chemostat with virus and virophage (purple triangles).

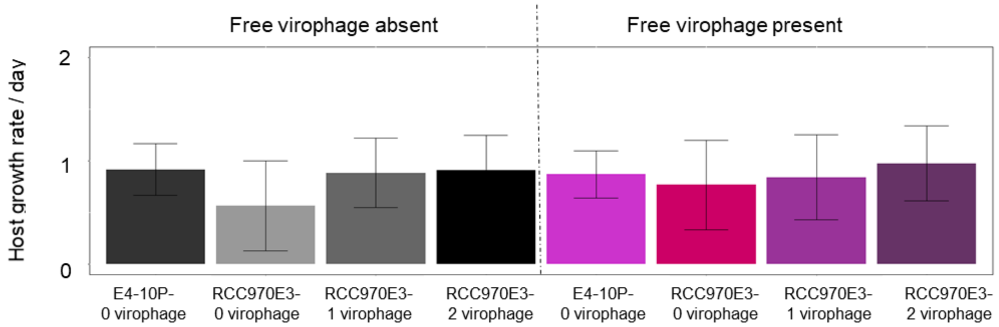


**Fig. S3.** Average host growth rate per day (5 days) in the absence and presence of free virophage for different host strains (mean ± s.d., n=5): Host strains vary in the number of virophage integrated: host strain E4-10P with no virophage integrated; and host-1virophage RCC970-E3 and host-2virophage RCC970-E3 are derived from host strain RCC970-E3 having 1, 2 and 0 virophage copies integrated in their genome. There were not significant differences in host growth in the presence or absence of free virophage neither depending on the number of integrated copies. Therefore, neither free nor 1-2 virophage integrations cause a cost for host growth under our experimental conditions.
